# Widespread Receptive Field Remapping in Early Visual Cortex

**DOI:** 10.1101/2023.05.01.539001

**Authors:** Sachira Denagamage, Mitchell P. Morton, Nyomi V. Hudson, Anirvan S. Nandy

**Affiliations:** Department of Neuroscience, Yale University, New Haven; Interdepartmental Neuroscience Program, Yale University, New Haven; Kavli Institute for Neuroscience, Yale University, New Haven; Wu Tsai Institute, Yale University, New Haven; Department of Psychology, Yale University, New Haven

## Abstract

Our eyes are in constant motion, yet we perceive the visual world as stable. Predictive remapping of receptive fields is thought to be one of the critical mechanisms for enforcing perceptual stability during eye movements. While receptive field remapping has been identified in several cortical areas, the spatiotemporal dynamics of remapping, and its consequences on the tuning properties of neurons, remain poorly understood. Here, we tracked remapping receptive fields in hundreds of neurons from visual Area V2 while subjects performed a cued saccade task. We found that remapping was far more widespread in Area V2 than previously reported and can be found in neurons from all recorded cortical layers and cell types. Surprisingly, neurons undergoing remapping exhibit sensitivity to two punctate locations in visual space. Furthermore, we found that feature selectivity is not only maintained during remapping but transiently increases due to untuned suppression. Taken together, these results shed light on the spatiotemporal dynamics of remapping and its ubiquitous prevalence in the early visual cortex, and force us to revise current models of perceptual stability.

## INTRODUCTION

Our early visual system is wired to store information in eye-centered (retinotopic) coordinates. With each movement of the eyes, the image falling onto the retina shifts rapidly, as does the visual information arriving at neurons in the cortex. Despite this, our perception of the world remains seamless and stable, implying that the visual system is able to compensate for self-generated movements. Previous research has suggested that receptive field (RF) remapping could contribute to this stability^1,2^. Remapping refers to the phenomenon in which neurons transiently shift their locus of spatial sensitivity (i.e. receptive field) before the onset of a saccadic eye movement towards their future, post-saccadic location. Remapping is considered to be a predictive mechanism because it both precedes and is temporally locked to eye movement initiation, and therefore requires advance information about both the timing and trajectory of an upcoming saccade. This information is thought to be conveyed through a corollary discharge signal originating in the brain regions responsible for initiating eye movements^1,3,4^.

Receptive field remapping has been reported in many visual areas, including V1^5^, V2^5^, V3^5^, V3A^5^, V4^6,7^, LIP^2^, FEF^8,9^, and SC^10–12^. In the cortex, early visual areas such as V1 and V2 are thought to have a low proportion of neurons that exhibit remapping, with higher order visual areas having a greater proportion^5^. Remapping has also been observed in humans using functional magnetic resonance imaging (fMRI)^13–15^ and electroencephalography (EEG)^16–19^, and an array of studies have reported on the behavioral consequences of remapping. Recently, it has also become clear that there may be multiple forms of remapping^6,7^, with the exact mode of remapping potentially depending on the brain region or cell type studied, or task demands. On one hand, remapping to the future receptive field (‘forward remapping’; **Figure 1A**) is thought to link pre- and post-saccadic representations of visual space, thus helping to maintain perceptual continuity and stability^2,5^. By contrast, remapping towards the saccade target (‘convergent remapping’; **Figure 1B**) may serve to transiently enhance processing of visual information near that target^7,20,21^. More recent evidence has suggested that both forms of remapping may exist within the same neurons^7^. There also remains concern that inconsistencies across some of these studies, which used different experimental paradigms in different visual areas, could have contributed to these divergent findings^22^. Furthermore, while work in human psychophysics has confirmed that remapping preserves some^17,23^ but perhaps not all^24^ visual feature selectivity at the level of perception, whether feature selectivity is preserved in individual neurons as they remap remains an open question. Indeed, despite extensive study, much remains unknown about the properties and extent of receptive field remapping, in large part due to limitations of the spatiotemporal resolution at which the phenomenon was studied^25–28^.

**Figure 1.**
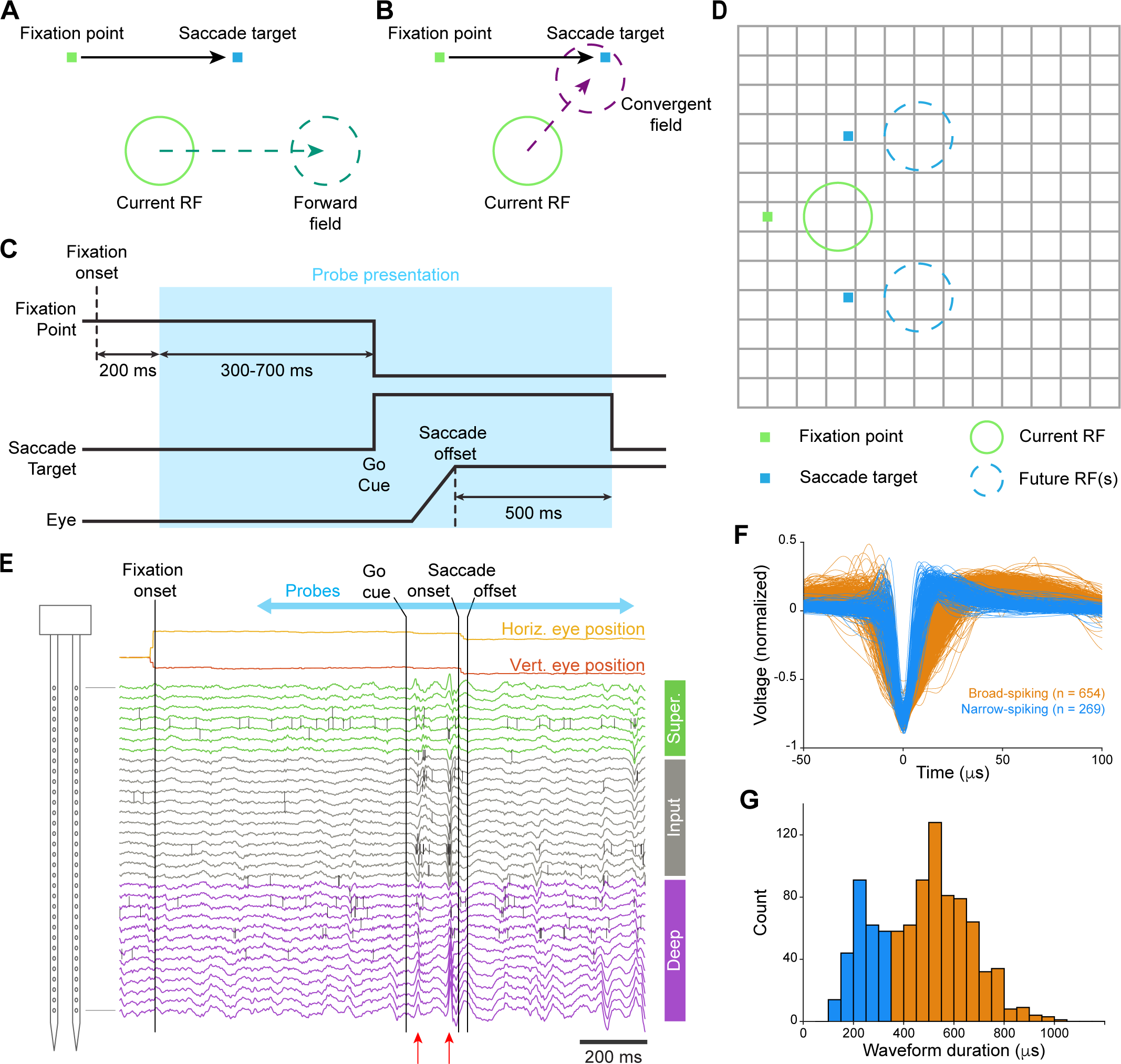
Task Design and Single Unit Recordings. **(A)** Forward remapping shifts the current receptive field by the vector of the upcoming saccade to form a forward field. **(B)** Convergent remapping shifts the current receptive field towards the saccade target to form a convergent field. **(C)** Progression of a trial during the cued saccade task as the subject holds fixation, executes a saccade to the target, and then fixates on the target. The period in blue indicates probe presentation. A reward is delivered after fixating on the target for 500 ms. **(D)** Fixation point, saccade targets, and stimulus grid layout during the cued saccade task. **(E)** Snippet of data from one probe shank during a trial of the cued saccade task. LFP traces are color by cortical layer, and spikes are overlaid on their channel of origin as vertical lines. Red arrows indicate synchronized spiking and local field potential deflections along the depth of the cortex in response to a stimulus being flashed in the receptive field of the recording site. **(F)** Action potential waveforms of all 923 single units. Units were classified as either narrow-spiking (blue) or broad-spiking (orange) on the basis of their peak-to-trough waveform duration. **(G)** Distribution of waveform durations for single units from (F). The distribution shows bimodality for the two unit types (Hartigan’s dip test; p = 1.15*10^-4^).

Here, we examined receptive field remapping in an early visual area (Area V2) with high-density electrode arrays and a stimulation paradigm that provided significantly improved spatiotemporal resolution. With this approach, we were able to track the time course of remapping in discrete neural subpopulations in the laminar cortical circuit, allowing us to test whether remapping is a global, trans-laminar phenomenon, or restricted to a particular cortical layer or cell type. The use of oriented Gabor stimuli also allowed us to examine whether tuning for visual features is altered during remapping. We found that remapping was far more prevalent in Area V2 than previously thought, and that it occurred in all recorded subpopulations. Neurons exhibit transient sensitivity to two punctate locations in visual space during remapping, similar to activity patterns found in the frontal cortex^29^ but unlike that reported in Area LIP^30^. Furthermore, peri-saccadic firing rate suppression results in a transient increase in orientation selectivity during remapping.

## RESULTS

We designed a cued saccade task, in which subjects held fixation for a variable delay period prior to initiating a saccade in response to a target point appearing in the periphery (**Figure 1C-D**). The simultaneous disappearance of the fixation point served as the go cue. After executing an accurate saccade, subjects then had to continue holding fixation at the target point to receive a reward. To prevent subjects from preemptively planning a saccade prior to the go cue, both the saccade target location and the delay period duration were pseudo-randomized. The target location was drawn from one of two possible locations, while the delay period duration was drawn from an exponential distribution. While the subjects executed these eye movements, oriented Gabor stimuli were continuously presented on a 13 x 13 grid spanning the visual region of interest at 60 Hz. On each frame of stimulus presentation, a single stimulus drawn from one of 6 random orientations was presented at a single grid location. Two rhesus macaques were trained to perform this task, and demonstrated consistent performance across trials and days (**Figure S1**).

While subjects performed the cued saccade task, we recorded neural activity from well-isolated single units in Area V2 using linear array electrodes (**Figure 1E**). The use of an artificial dura (**Figure S2A**) allowed us to clearly visualize the cortical vasculature, which provided landmarks for tracking successive probe insertion sites (**Figure S2B**). We confirmed that individual electrode penetrations were perpendicular to the cortical surface, and therefore in good alignment with individual cortical columns, by mapping receptive fields along the depth of the cortex (**Figure S2C**). Laminar boundaries were identified with current source density (CSD) analysis^31,32^ (**Figure S2D**), and units were classified as belonging to either the superficial (II/III), input (IV), or deep (V/VI) layers. Single units were also identified as either narrow- or broad-spiking based on their waveform duration (**Figure 1F-G**). In total, 923 single units were recorded, 822 of which were significantly visually responsive and included in our subsequent analyses.

### Widespread Pre-Saccadic Remapping in Area V2

For each recorded single unit, we computed spatial sensitivity maps as a function of time relative to saccade onset (**Figure 2A; Figure S3A-C; Video S1 and S2**). We found that 73% of V2 units showed pre-saccadic remapping to the future receptive field (forward remapping) before the start of an eye movement. We found no evidence of widespread remapping towards the saccade target (convergent remapping), suggesting that this phenomenon may only arise in higher order visual areas. Using these spatial maps, we calculated the relative spatial distribution of sensitivity between the current and future receptive fields as a function of time (**Figure 2B; see Figure S3D for raw firing rate traces**). These time courses show that the handoff of spatial sensitivity begins well before saccade onset and has largely completed by the time the eyes begin to move. Surprisingly, our results suggest that for this brief period, these units are significantly responsive to two discrete locations in visual space. This pattern of spatial sensitivity transfer is not unique to any of the neural subpopulations that were recorded, and instead occurs in all three layers as well as in both broad- and narrow-spiking units (**Figure 2C**). Indeed, the timing of this transfer is also consistent across subpopulations, initiating approximately 40 ms before the saccade (**Figure 2D**), after spike binning normalizes for feed-forward signaling delays (**Figure S3C**).

**Figure 2.**
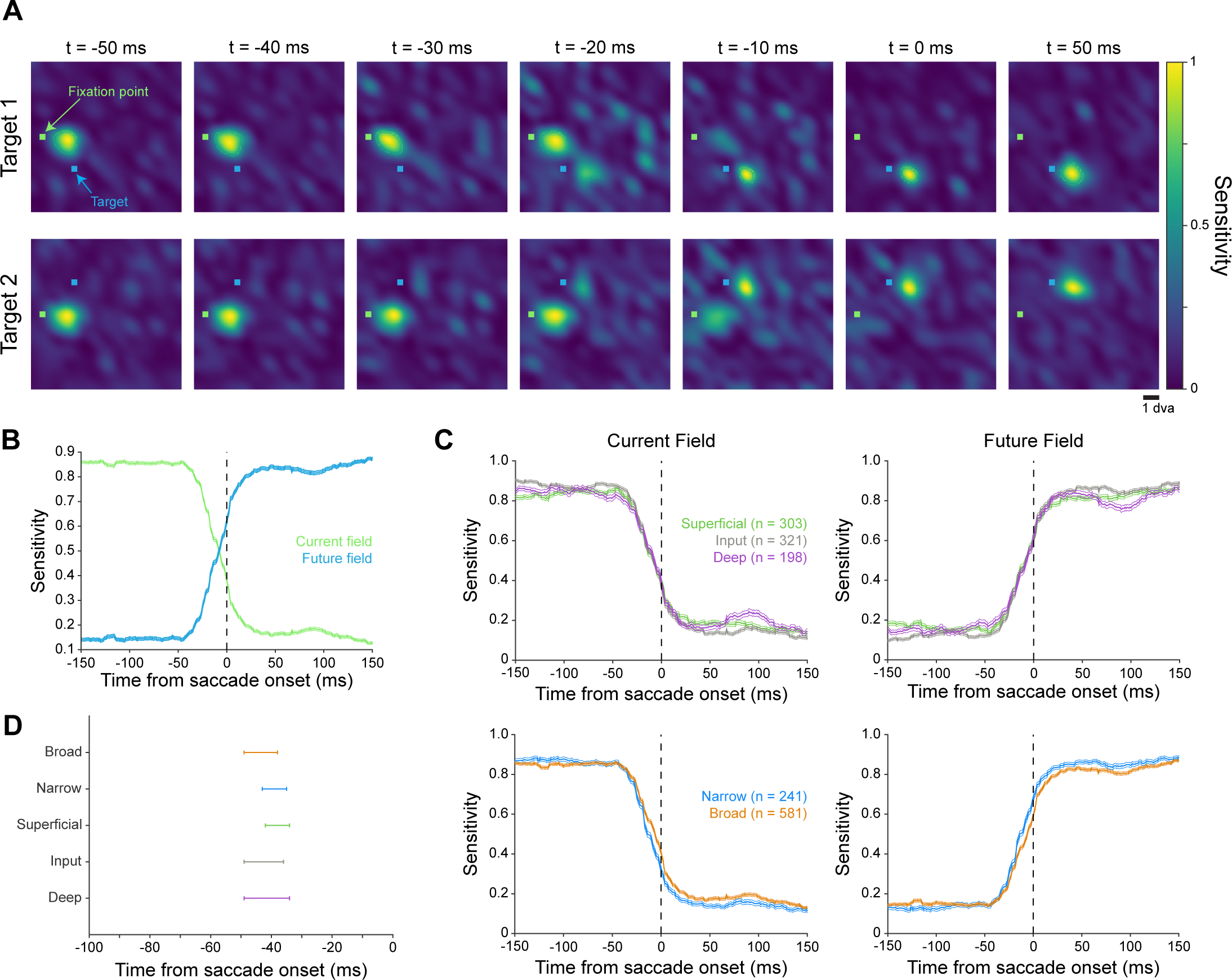
Receptive Field Remapping is Widespread in Area V2. **(A)** Receptive field location for an example single unit during remapping. Times are relative to saccade onset. Each row shows remapping during saccades to one of the two saccade targets. Fixation point and saccade target are overlaid in green and blue, respectively. Heatmaps at each timepoint are individually normalized to account for possible changes in firing rate. **(B)** Normalized sensitivity at the current and future field locations for all single units (n = 822), as a function of time relative to saccade onset. Error bars indicate standard error of the mean. **(C)** Normalized sensitivity at the current and future field locations as a function of time relative to saccade onset for each of the recorded neural subpopulations. Error bars indicate standard error of the mean. **(D)** The time relative to saccade onset at which remapping initiates for each of the recorded neural subpopulations. Error bars indicate bootstrapped 95% confidence intervals.

### Tracking Receptive Field Trajectories in Principal Component Space

To determine whether the transition in spatial activity patterns could be identified without pre-defined coordinates for the current and future fields, we performed unbiased dimensionality reduction analyses (principal component analysis, PCA) on our data. We vectorized the 13 x 13 stimulus grid into a 169-dimensional space with neural response values that changed over time for each unit; each timepoint was a single sample for the PCA. As all of the units from each session had largely overlapping receptive fields, and thus similar activity patterns in this space, they were fed into the PCA together. This approach produces a low-dimensional representation of receptive field trajectories during remapping (**Figure 3A; Figure S4A**) by, in effect, detecting the features of the spatial distribution of sensitivity (see Figure 2A for an example) that account for the greatest variance across time. Strikingly, the prominent features of this analysis are consistent across sessions, animals, and cortical layers, with a characteristic V-shaped trajectory (**Figure S4B; Figure S5; Figure S6**). These results can also be normalized and pooled for units across all sessions to produce the receptive field trajectory of an average V2 neuron during remapping (**Figure 3B**), which again illustrates this characteristic shape. To determine which features of the neural activity were being identified by the PCA, we compared values along the 1st PC dimension as a function of time with our previous current/future field sensitivity traces. We found that the 1st PC dimension closely tracks the relative distribution of spatial sensitivity between the current and future field (**Figure 3C**; Pearson’s correlation = 0.996), remaining relatively stable until shortly before saccade onset and then shifting to values at the other extreme. Further examination of the 1st PC time courses revealed that this transition was consistent across all unit subpopulations (**Figure 3D-E**) and was initiated at a time that closely matches the results from our spatial map analysis (**Figure 3F**). The 2^nd^ PC, on the other hand, appears to track the percentage of the total response that is contained within the receptive field relative to the baseline response at non-receptive field positions (**Figure S4C**), which is an indirect measure of changes in peri-saccadic firing rate (**Figure S4D**) presumably caused by saccadic suppression. Therefore, it is the temporally overlapping effects of receptive field remapping and saccadic suppression that generate V-shaped trajectories in principal component space. Most notably, even with this approach that makes no assumptions about particular patterns of spatial sensitivity at particular moments in time, translational shifts to the future field remain the dominant feature of remapping in Area V2.

**Figure 3.**
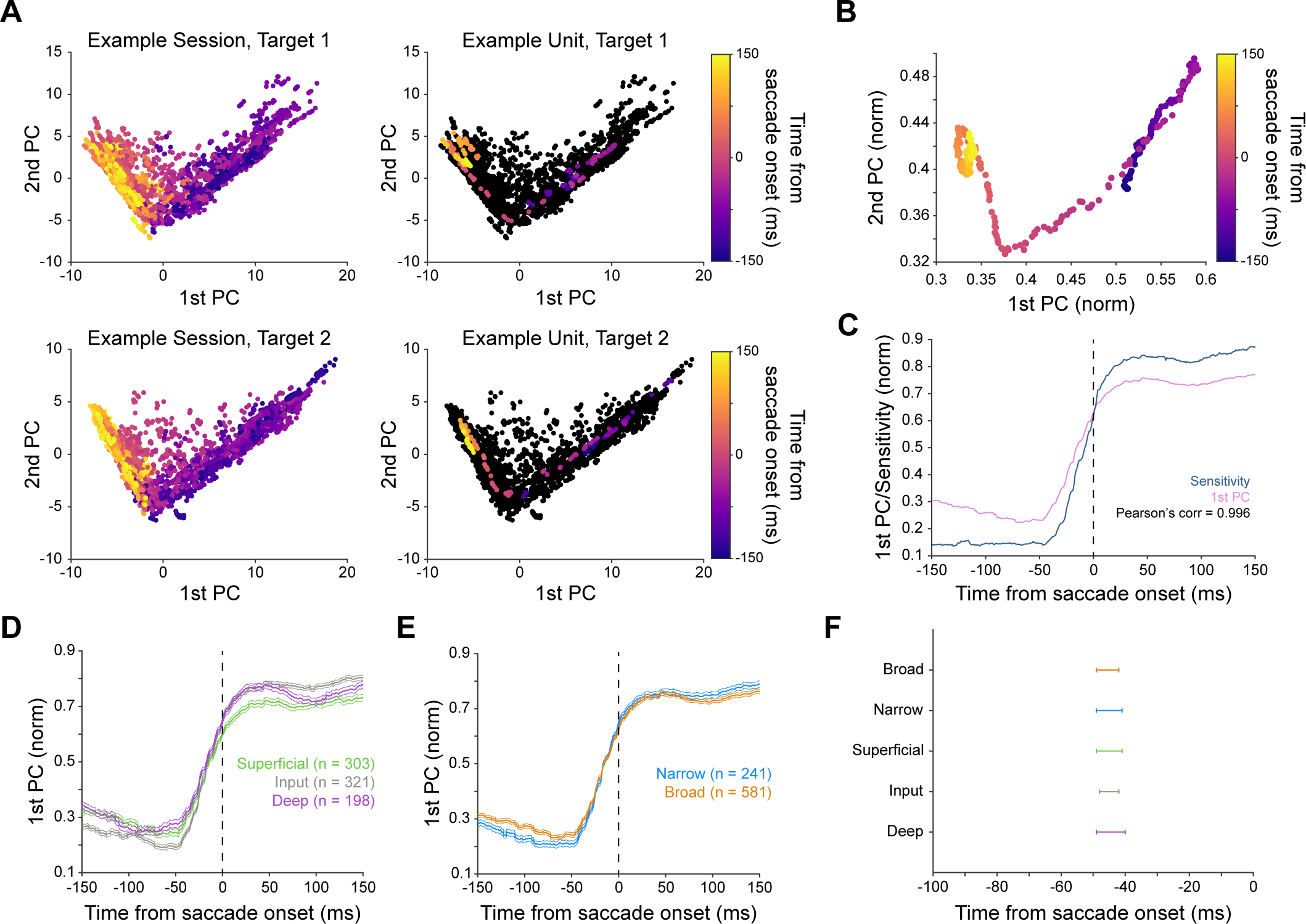
Tracking Remapping Trajectories in Principal Component Space. **(A)** Left, principle component trajectories of all single units from an example session. Right, principle component trajectory of an example single unit from the same session. Top, target 1. Bottom, target 2. Each point represents a single timepoint from a single unit. On average across sessions and targets, PC1 explains 11.3% of the variance, and PC2 explains 3.3% of the variance. **(B)** Averaged remapping trajectory in principal component space across all single units for both targets. **(C)** Correlation between values along the 1^st^ principle component axis and sensitivity at the future field (Figure 2B). Pearson’s correlation = 0.996. **(D)** Time course of values along the 1^st^ principle component axis for all recorded layers. Error bars indicate standard error of the mean. **(E)** Time course of values along the 1^st^ principle component axis for both recorded unit types. Error bars indicate standard error of the mean. **(F)** The time relative to saccade onset at which remapping initiates for each of the recorded neural subpopulations based on the 1^st^ PC timecourses. Error bars indicate bootstrapped 95% confidence intervals.

### Firing Rate Suppression Drives a Transient Enhancement of Tuning During Remapping

To determine whether stimulus selectivity may be altered during remapping, we generated tuning curves for each unit during pre-saccadic, saccade planning, and post-saccadic periods (**Figure 4A**). The remapping period was defined as -75 to -25 ms relative to saccade onset to cover the transition between the current and future field, while the pre- and post-saccadic periods were well before and after saccade onset respectively. From these tuning curves, we computed the preferred orientation of each unit, as well as their orientation selectivity index (OSI; **Figure 4B-C**) and circular variance (**Figure 4D-E**). A higher OSI indicates a greater preference for one orientation over the orthogonal orientation. A circular variance of 1 would reflect equal responses to all orientations, while a circular variance of 0 would reflect responsiveness to only a single orientation. When comparing OSI and circular variance across the three conditions, we found that orientation tuning was transiently increased during saccade planning before returning to baseline levels. We next asked whether this increase in tuning could be the result of a firing rate change. Population averaged tuning curves revealed that the firing response to stimuli at both the preferred and non-preferred (orthogonal) orientations were suppressed during remapping (**Figure 4F-G**). Fitting the data from each unit with a Gaussian tuning curve also showed the same pattern of suppression (**Figure 4H**). Quantifying the half width at half height (HWHH) from the fitted curves revealed no changes in the overall shape of the tuning curves (**Figure 4I**). Thus, this change in tuning is largely driven by untuned suppression, as firing in response to both preferred and non-preferred orientations is suppressed, and not by divisive changes to the shape of the tuning curve, as the HWHH of the tuning curves remains unchanged.

**Figure 4.**
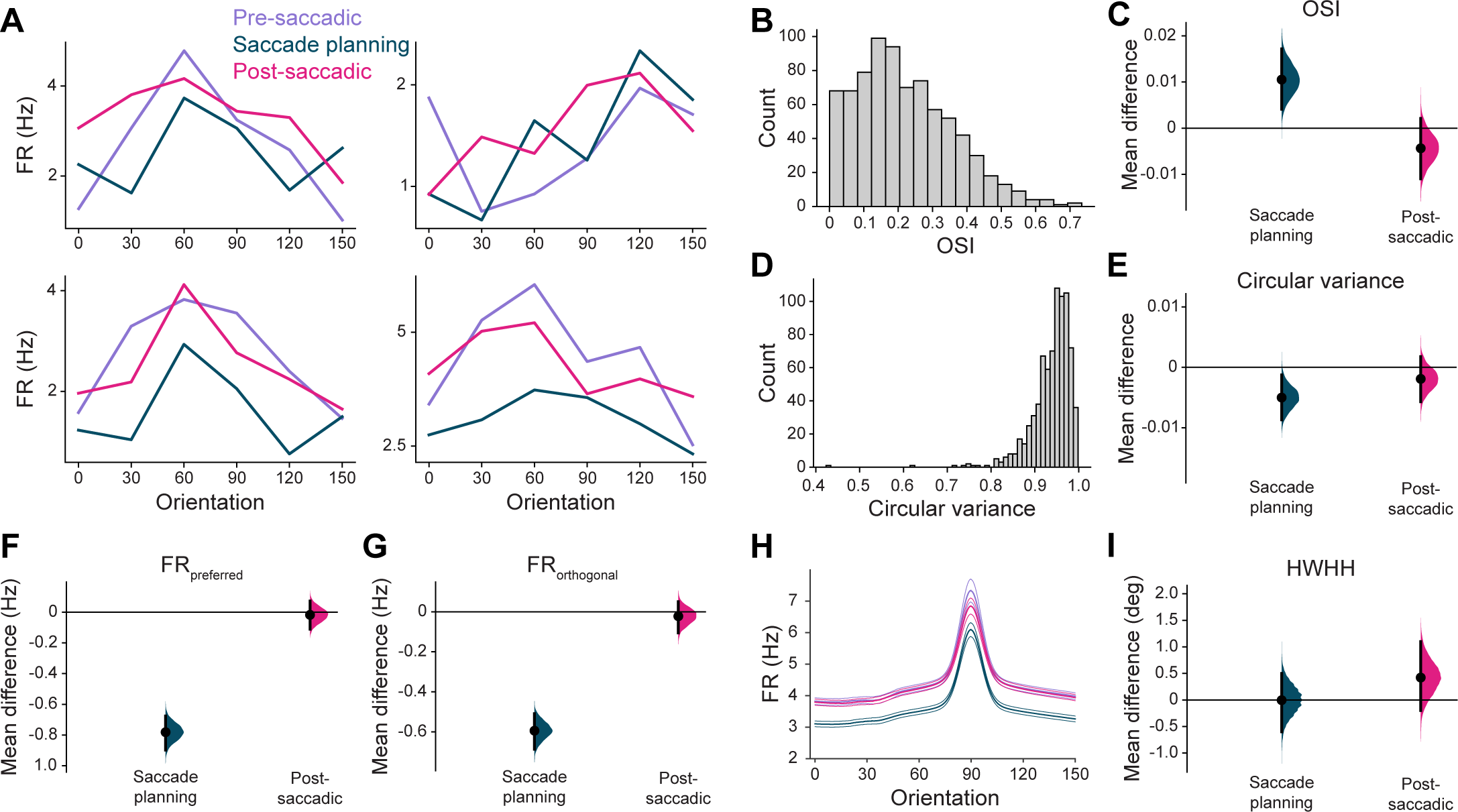
Orientation Selectivity is Transiently Increased During Saccade Planning. **(A)** Orientation tuning curves from four example single units during the pre-saccadic (-250 to -200 ms), saccade planning (-75 to -25 ms), and post-saccadic (150 to 200 ms) time periods. Times are relative to saccade onset. FR = firing rate. **(B)** Distribution of orientation selectivity index (OSI) for all single units during the pre-saccadic period. **(C)** Change in OSI during the saccade planning and post-saccadic periods, as compared to the pre-saccadic period. Error bars indicate bootstrapped 95% confidence intervals. **(D)** Distribution of circular variance for all single units during the pre-saccadic period. **(E)** Change in circular variance during the saccade planning and post-saccadic periods, as compared to the pre-saccadic period. Error bars indicate bootstrapped 95% confidence intervals. **(F)** Change in firing rate in response to presentation of a stimulus at the preferred orientation during the saccade planning and post-saccadic periods, as compared to the pre-saccadic period. Error bars indicate bootstrapped 95% confidence intervals. **(G)** Change in firing rate in response to presentation of a stimulus at the non-preferred (orthogonal) orientation during the saccade planning and post-saccadic periods, as compared to the pre-saccadic period. Error bars indicate bootstrapped 95% confidence intervals. **(H)** Average of Gaussian tuning curve fits across all single units during the pre-saccadic, saccade planning, and post-saccadic time periods. Error bars indicate standard error of the mean. **(I)** Change in half width at half height (HWHH) of the fitted tunning curves during the saccade planning and post-saccadic periods, as compared to the pre-saccadic period. Error bars indicate bootstrapped 95% confidence intervals.

## DISCUSSION

We used high density receptive field mapping during a cued saccade task in combination with laminar electrophysiology to study receptive field remapping in Area V2. We found that remapping was widespread in Area V2, and was present across all recorded cortical layers (superficial, input, deep) and unit types (narrow- and broad-spiking). We identified forward remapping towards the future receptive field location as the dominant mode of remapping in Area V2. We demonstrated that these findings were not contingent on any assumptions about receptive field structure or direction of remapping, as an unbiased dimensionality reduction approach arrives at the same result. Further, we tested for changes in tuning during saccade planning and execution, and found that a brief suppression of firing rates drives transiently increased tuning during saccade planning.

Our data demonstrate that receptive field remapping is far more widespread in the early visual cortex than was previously appreciated. Prior work has suggested that the proportion of neurons that underwent remapping increased progressively along the visual stream, with early visual cortex, including areas such as V1 and V2, having relatively few neurons that remapped^5^. It was thought that this low proportion may have reflected the prevalence of remapping in a particular cell type, and the absence of remapping in others. However, our results indicate that remapping is a global phenomenon that occurs in a majority of neurons in Area V2 and is not restricted to a particular cell class or layer. This discrepancy can likely be explained by differences in experimental approach. The only prior electrophysiological study to test for remapping in Area V2^5^ presented stimuli at just four timepoints relative to a go cue, resulting in a relatively course temporal sampling. Given technical limitations at the time, the neural responses to these stimuli were not aligned to saccade onset, and trial-to-trial variability in behavioral response times may therefore have also limited the temporal clarity of the data. Flashing of single probes at discrete timepoints may also draw attention to the probe, resulting in attentional remapping that may affect visual responses^18^. In our study, we are able to generate continuous time courses of spatial sensitivity at both the current and future receptive field that are aligned to the onset of a saccade, and with this approach we observe a much higher proportion of neurons undergoing remapping.

The functional role of remapping remains a largely open question, although the prevailing hypothesis is that remapping plays a significant role in maintaining perceptual stability across eye movements^1,2^. Forward remapping towards the future receptive field may allow for visual processing before the saccade to occur in a post-saccadic frame of reference, thereby allowing the visual system to maintain a spatial frame of reference that would otherwise be disrupted by an eye movement. Given the existence of multiple modes of remapping^6,7^ and presumed differences in their relative prevalence across cortical areas, it is also possible that the functional significance of remapping varies from region to region. Given the shifts in receptive field location, as well as the potential for spatial sensitivity at split locations in space, remapping may also be responsible for the mis-localization of stimuli around the time of saccades^33–35^.

Given the prevalence of remapping in Area V2, it is also worth considering whether remapping may be important in other early visual cortical areas, such as Area V1. Remapping is often considered to be a result of, or at least a correlate to, the allocation of spatial attention^18,19,36,37^. However, V1 and V2 serve different roles in attentional allocation, with V1 generating a saliency map that guides attention^38–42^, while V2 is not known to be significantly involved. Thus, remapping may be an undesirable property in the V1 neural population that conflicts with its role in attention guidance, resulting in very few V1 neurons showing remapping. Nonetheless, further studies are needed to identify possible differences in remapping along the visual hierarchy.

Corollary discharge signaling from the superior colliculus through the thalamus is thought to be responsible for initiating receptive field remapping in the cortex. Pharmacological interventions have demonstrated that thalamic inactivation impairs performance in a double-step saccade task^4^ and limits cortical remapping^29^, confirming the role of the thalamus as a relay station for this signal. Recent evidence suggests that thalamic projections to the visual cortex may also be used for distinguishing between self-generated and saccade-generated visual motion in both primates^43^ and rodents^44^. Interestingly, there may also be a high degree of redundancy in pathways for the updating of peri-saccadic spatial information, as behavioral performance in a double-step saccade task was found to be impaired in split-brain monkeys, but could be recovered substantially with training^45^. Our results are consistent with a thalamic origin for the signal initiating receptive field remapping. Our timing analysis, which normalizes for feedforward timing delays as a byproduct of spike binning, found no significant differences in the onset of receptive field remapping across layers. This suggests that the feedforward transfer of visual information across layers, starting with the input layer and then spreading to the superficial and deep layers, also carries the remapping signal. Thus, the remapping signal appears to first arrive in the input layer of V2, which is where thalamic inputs to V2 terminate in both macaques and other primates^46–48^. Recent evidence from an intermediate visual region, Area V4, suggests that thalamic projections to the input layer may also be responsible for initiating saccadic suppression^43^, raising the possibility that a common signaling pathway from the thalamus may be responsible for both phenomena. An alternative possibility is that receptive field remapping in V2 is the result of feedforward input from V1. However, we consider this to be the less likely option given the importance of V1 in maintaining an attentional saliency map, as discussed above, and the much stronger input that V2 receives from the pulvinar^47^, the thalamic nucleus thought to relay saccade-related signals.

One understudied aspect of remapping has been the question of whether feature selectivity remaps alongside the spatial receptive field. This question is of critical behavioral importance, as perceptual stability requires the ability to identify stimuli as well as to locate them. Thus far, research on this topic has been both sparse and conflicting. In LIP, it is thought that stimulus tuning is preserved during remapping^49^, while in MT, there is no evidence of tuning in remapped fields^50^. Here, we show that one prominent aspect of feature encoding in Area V2, orientation tuning, persists during remapping. Indeed, we find that orientation selectivity transiently increases due to the suppression of overall firing during the saccade planning period.

Together, our results reveal the widespread nature of receptive field remapping in the early visual cortex and suggest that the fundamental computations underlying perceptual stability are enacted from these early stages. Furthermore, we demonstrate that remapping overlaps and interacts with changes in feature tuning that are driven by saccadic suppression of neural firing. The cortical column-wide nature of the changes suggests that remapping is conveyed as a global signal to the early visual cortex. The fact that neurons exhibit transient split sensitivity to two punctate locations in space necessitates a rethinking of the nature and functional role of remapping. Further experiments are needed to both fully characterize sensitivity and tuning changes at a sub-receptive field level and to elucidate the neural circuits that enable these phenomena.

### Limitations of the study

This study has several limitations that should be considered alongside our findings. For one, our approach treats the receptive field of V2 neurons as homogenous and is unable to resolve potential differences in remapping across subfields. Second, it is unknown whether neural response latencies may change during remapping, and we assume that they are constant when settings our binning windows (Figure S3A-B). And lastly, despite our approach providing us with cell type- and layer-specific insights into remapping, we remain limited by the tools available to us and are unable to causally link these effects to a specific signaling pathway.

## ACKNOWLEDGEMENTS

This research was supported by NIH/NEI R01 EY032555, NARSAD Young Investigator Grant, Ziegler Foundation Grant and Yale Orthwein Scholar Funds to ASN, NIH/NINDS training grants T32-NS007224 and T32-NS041228 to SD, and by an NIH/NEI core grant for vision research P30 EY026878 to Yale University. We would like to thank the veterinary and husbandry staff at Yale for excellent animal care. We would like to thank John Reynolds for helpful comments on the manuscript.

## AUTHOR CONTRIBUTIONS

SD & ASN conceptualized the project. SD, MPM, and NVH collected the data. SD analyzed the data. ASN supervised the project. SD & ASN wrote the manuscript.

## DECLARATION OF INTERESTS

The authors declare no competing interests.

## INCLUSION AND ETHICS

We support inclusive, diverse, and equitable conduct of research.

## Supplementary

**Figure S1.**
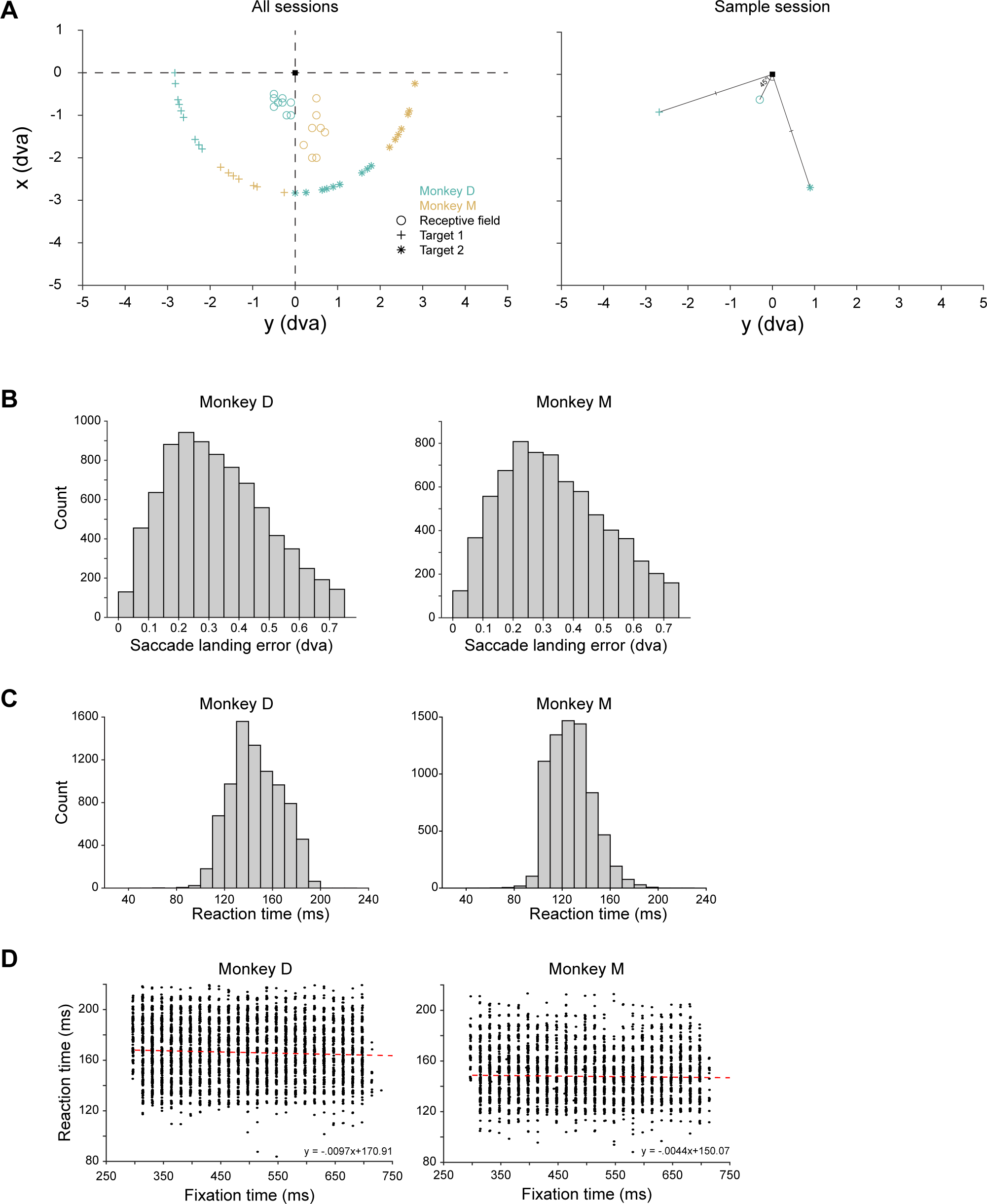
Behavioral Performance. **(A)** Receptive field and saccade target locations for all sessions (left) and one sample session (right). **(B)** Distribution of saccade landing errors for both subjects. Landing error was defined as the distance between the saccade target and the terminal point of the saccade. **(C)** Distribution of reaction times for both subjects. **(D)** Relationship between the trial-by-trial pre-saccadic fixation time (pseudorandomly determined for each trial) and reaction time. The dashed red line depicts a linear least-squares fit. The flat relationship between fixation time and reaction time suggests that subjects are not able to anticipate the timing of the go cue.

**Figure S2.**
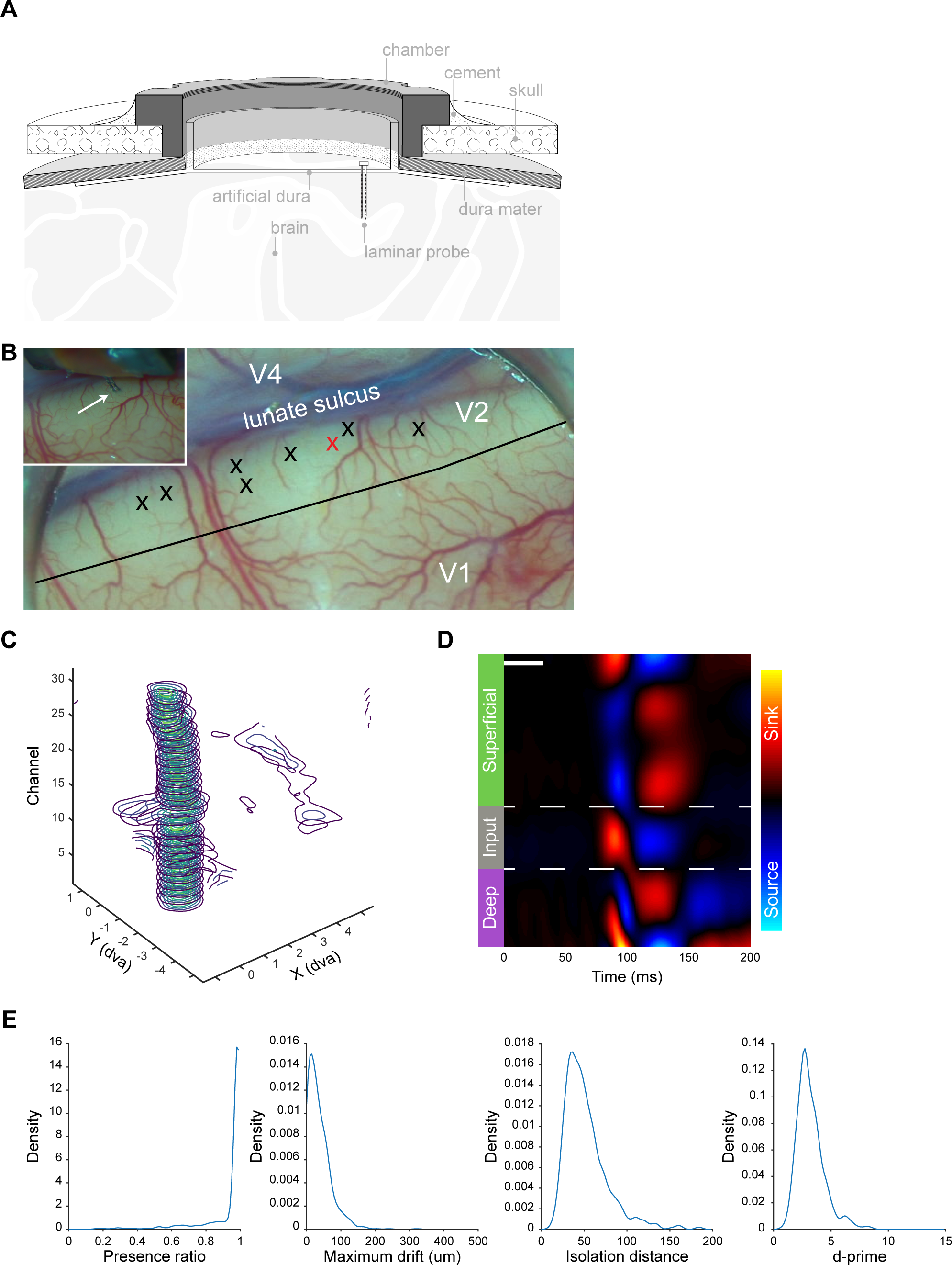
Recording Methodology. **(A)** Schematic of the artifical dura and recording chamber (see Methods). **(B)** Microscope image inside the recording chamber of one subject. Each ‘X’ indicates a cortical recording site. The inset shows a close-up image of the linear array probe penetration at the site marked in red. The black line is the estimated boundary between Area V1 and Area V2 based on the distinct change in surface vasculature density between the two areas. **(C)** Receptive fields along the depth of one shank as computed from local field potential deflections. dva = degrees of visual angle **(D)** Current source density (CSD) analysis of the shank from (C). Blue indicates a current source, while red indicates a current sink. Dashed lines indicates laminar boundaries, as determined from the CSD. White bar indicates duration of stimulus presentation. **(E)** Kernel density distribution estimates for several metrics^51–53^ that quantify recording stability and single unit quality. Presence ratio reflects the proportion of a session during which a unit was present and firing action potentials. Maximum drift is the distance between the highest and lowest channels on which a unit was detected during a session. Isolation distance and d-prime quantify the separation of spike waveform clusters in principal component space.

**Figure S3.**
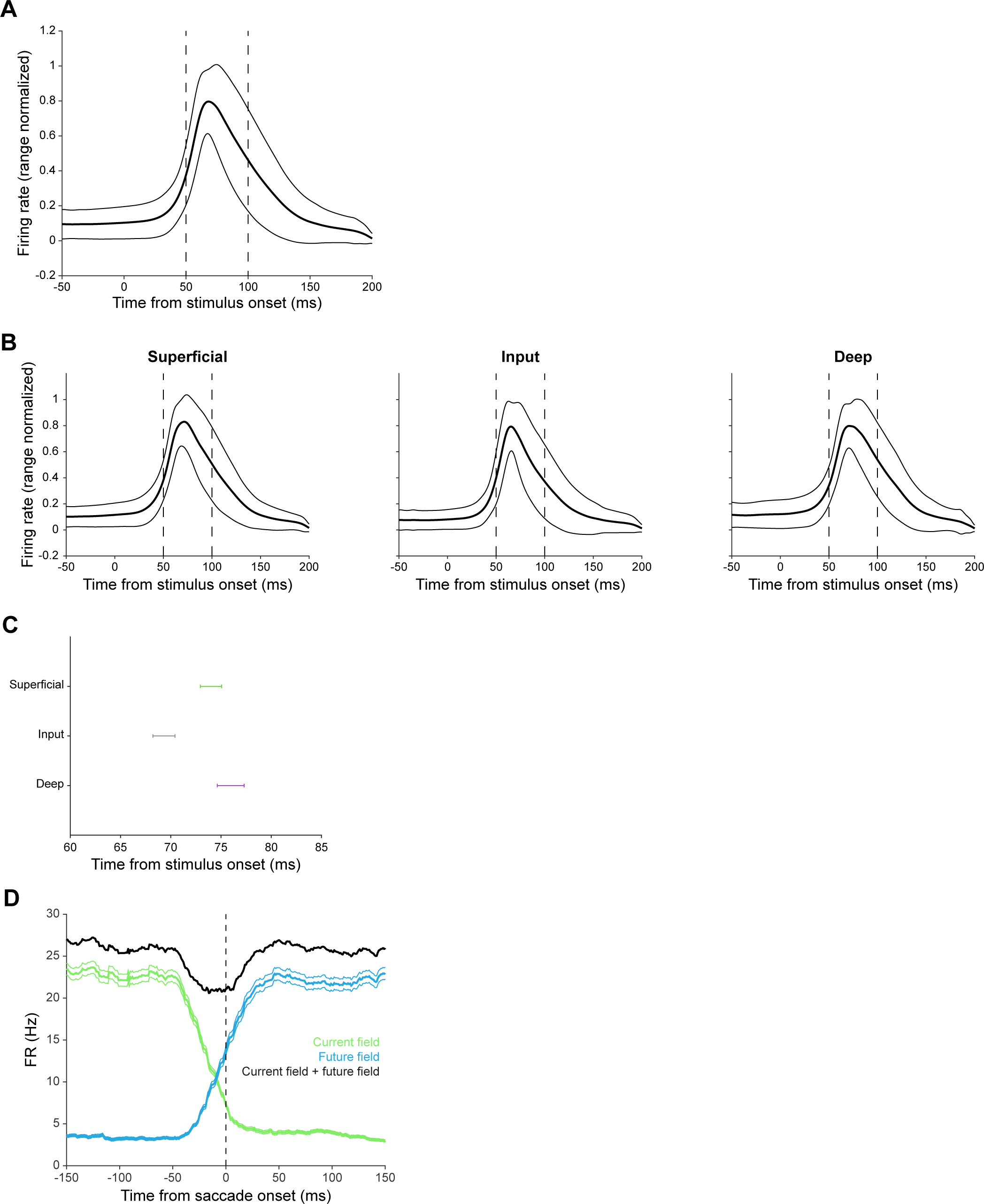
Stimulus-evoked firing rates at current and future fields. **(A)** Average firing response to stimulus flashes in the current receptive field during a baseline period before presentation of the go cue (n = 822 single units). Dashed vertical lines indicate the start and end of the binning window. Error bars indicate 95% bootstrapped confidence intervals. **(B)** Same as in (A), but separated by cortical layer (superficial, n = 303 single units; input, n = 321 single units; deep, n = 198 single units). Dashed vertical lines indicate the start and end of the binning window. Error bars indicate 95% bootstrapped confidence intervals. **(C)** Time of peak firing relative to stimulus onset for data in (B). Error bars represent bootstrapped 95% confidence intervals (superficial, n = 303 single units; input, n = 321 single units; deep, n = 198 single units). **(D)** Firing rate responses at current and future field locations (n = 822 single units). Contrast with sensitivity plots, which are normalized at each timepoint (Figure 2B). Error bars indicate standard error of the mean.

**Figure S4.**
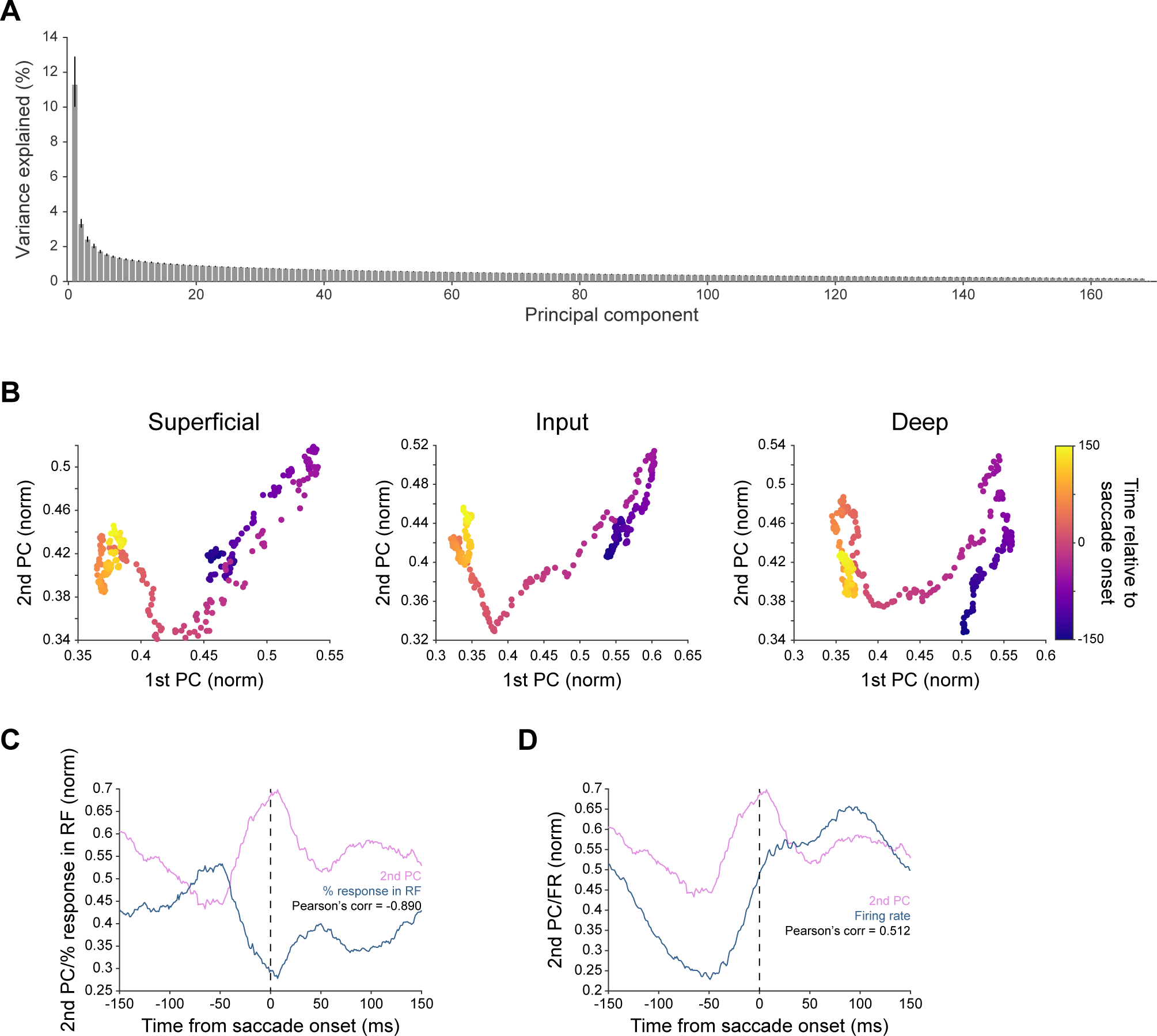
Remapping Trajectories are Tracked in Principal Component Space. **(A)** Variance explained by each principal component, averaged across sessions and targets. Error bars indicate bootstrapped 95% confidence intervals. **(B)** Averaged remapping trajectory in principal component space across all single units for both targets when PC analysis is performed on units from a given layer. **(C)** Correlation between values along the 2nd principal component axis and the percent of total response contained within the current and future fields (as opposed to firing evoked by stimuli landing outside the receptive field). Pearson’s correlation = -0.890. **(D)** Correlation between values along the 2nd principal component axis and firing rate. Pearson’s correlation = 0.512.

**Figure S5.**
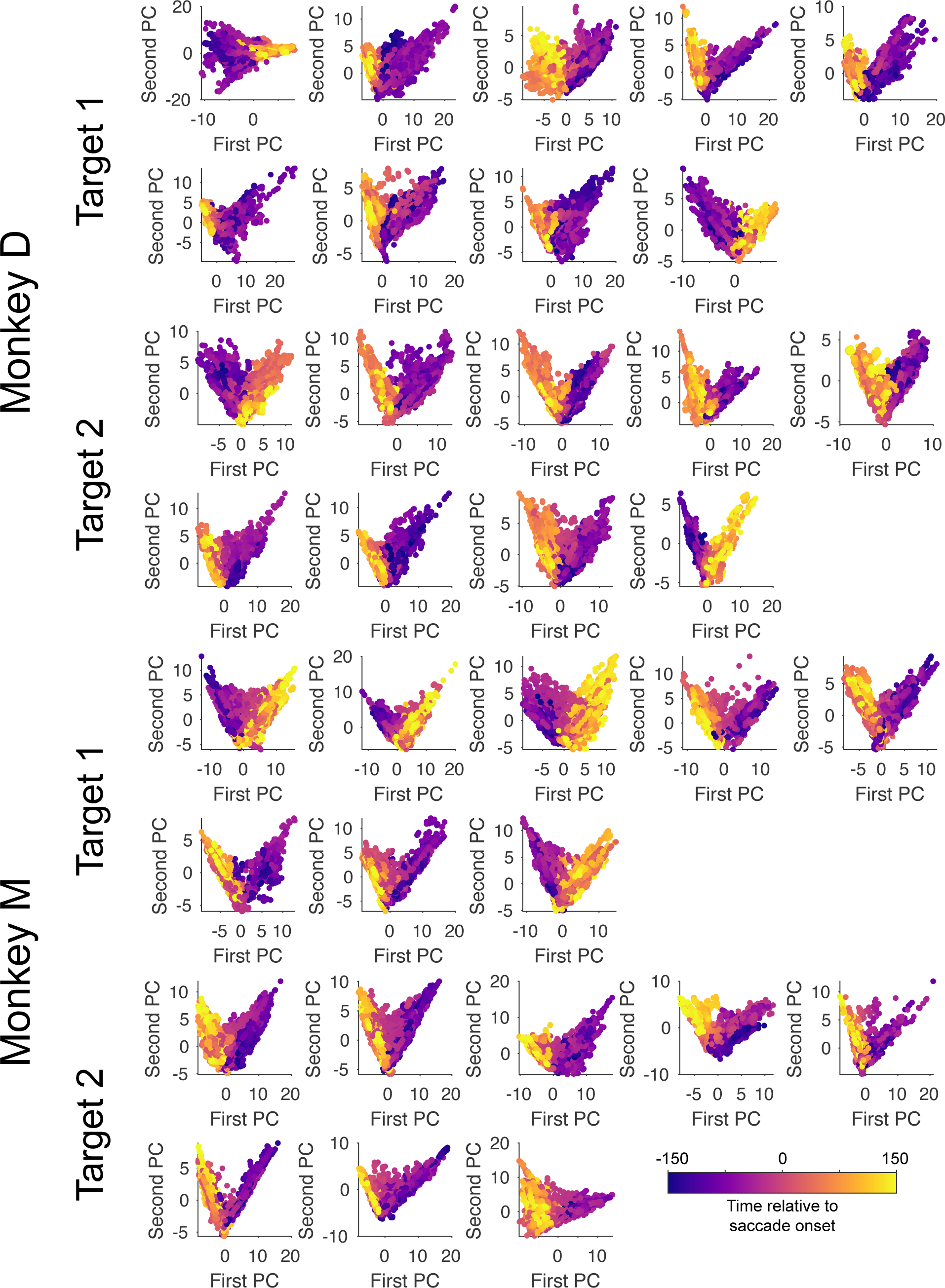
Principle Component Trajectories of All Sessions. Receptive field remapping trajectories in principal component space of all sessions. Across sessions, the data consistently shows a V-shaped trajectory in this space as remapping occurs.

**Figure S6.**
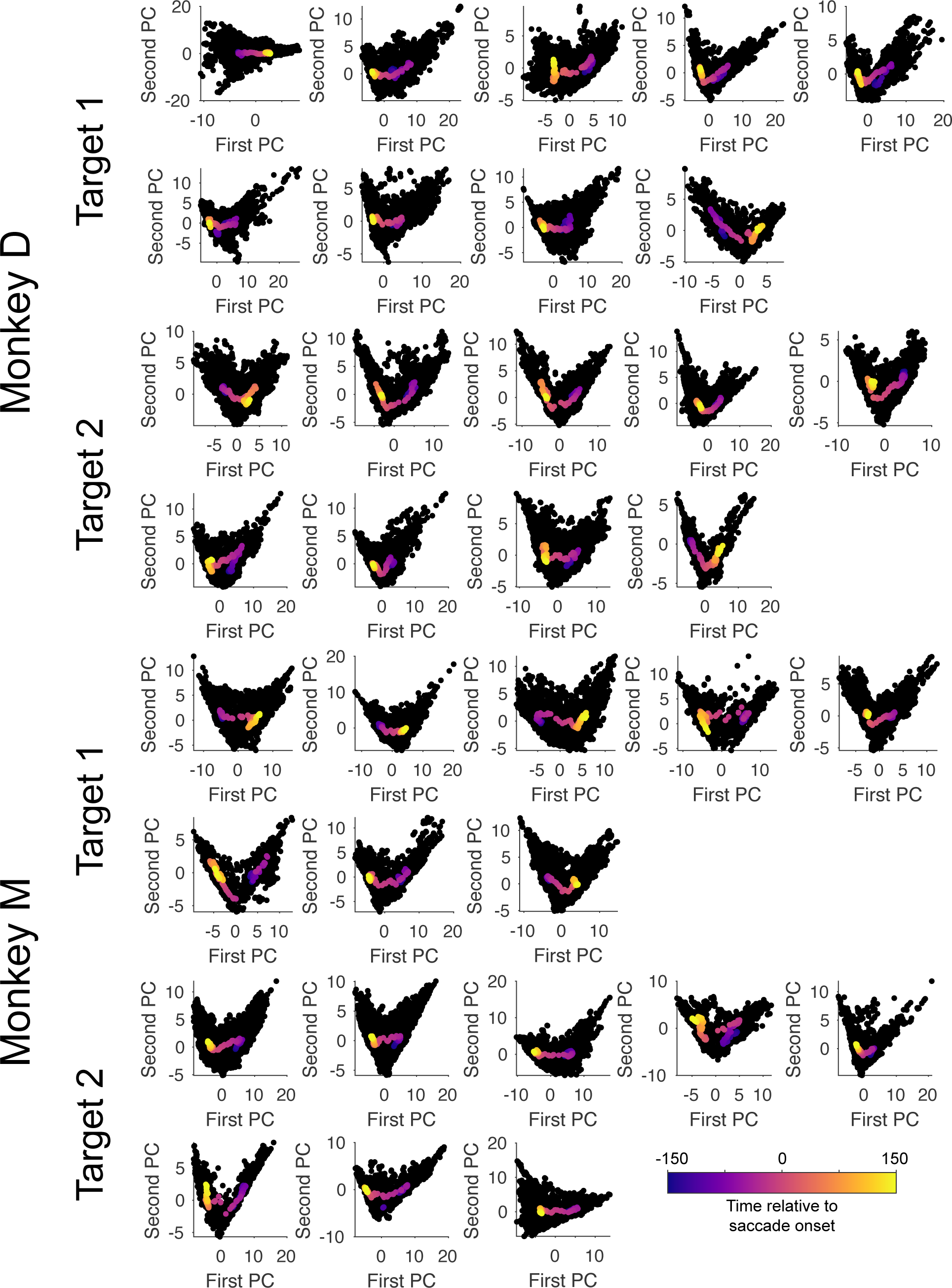
Average Principle Component Trajectories of All Sessions. Average receptive field remapping trajectories in principal component space of all sessions.

**Video S1. Continuous Tracking of Receptive Field Remapping**

Spatial location of an example unit’s receptive field sensitivity as a function of time (bottom left). Times are relative to saccade onset.

**Video S2. Continuous Tracking of Receptive Field Remapping**

Same as Video S1, but for another example unit from a different session.

## STAR METHODS

### KEY RESOURCES TABLE

| REAGENT or RESOURCE | SOURCE | IDENTIFIER |
| --- | --- | --- |
| **Experimental models: Organisms/strains** |  |  |
| Rhesus Macaques (*Macaca mulatta*) | Worldwide Primates | n/a |
| **Chemicals** |  |  |
| PEDOT:PSS | Sigma-Aldrich | 655201 |
| **Software and algorithms** |  |  |
| MATLAB | Mathworks | R2019a |
| Python | python.org | n/a |
| Kilosort2 | Pachitariu M & MouseLand | https://github.com/MouseLand/Kilosort |
| phy (spike curation GUI) | Rossant C & cortex-lab | https://github.com/cortex-lab/phy |
| Custom code and analyses | This paper | https://doi.org/10.5281/zenodo.11044507 |
| **Other** |  |  |
| Silicon probes | NeuroNexus | a2x32_6mm35_200_177 |
| RHD 512 channel recording controller | Intan | C3004 |
| 64 channel recording headstages | Intan | C3315 |
| nanoZ | White Matter LLC | n/a |

## RESOURCE AVAILABILITY

### Lead contact

Further information and requests for resources and reagents should be directed to and will be fulfilled by the lead contact, Sachira Denagamage.

### Materials availability

This study did not generate new unique reagents or materials.

### Data and code availability

All data reported in this paper will be shared by the lead contact upon request. All original code has been deposited at Zenodo and are publicly available as of the date of publication. DOIs are listed in the key resources table. Any additional information required to reanalyze the data reported in this paper is available from the lead contact upon request.

## EXPERIMENTAL MODEL AND SUBJECT DETAILS

Two male rhesus macaques (*Macaca mulatta,* D: age 6, M: age 8) were used as subjects in this study. All experimental procedures were approved by the Institutional Animal Care and Use Committee at Yale University, and conformed to NIH guidelines.

## METHOD DETAILS

### Experimental design

The study did not involve randomization or blinding, and we did not estimate sample-size before carrying out the study. No subjects or data were excluded from the study.

### Surgical procedures

Surgical procedures were similar to those described previously ^32,54,55^. Low-profile titanium recording chambers were implanted in two rhesus macaques. Using preoperative structural MRI and sulcal reconstruction, the chambers were targeted over the lunate sulcus, allowing access to Area V2 (left hemisphere in monkey M, right hemisphere in monkey D). The native dura mater overlying this region was removed and replaced with a transparent silicone artificial dura (AD). The AD allowed for visualization of area V2 and facilitated the targeting of electrode arrays.

### Electrophysiology

Prior to recording, 64-channel electrode arrays (‘laminar probes’; NeuroNexus Technologies, Inc.; 2 shanks, 32 channels/shank, 70 µm site spacing, 200 µm shank spacing) were electroplated (nanoZ, White Matter LLC) in a solution of poly(3,4-ethylenedioxythiophene) polystyrene sulfonate (*PEDOT*:PSS). At the start of each recording session, a laminar probe was lowered into Area V2 through the use of electronic micromanipulator (Narishige Inc.). Visual inspection of the cortical surface through a surgical microscope (Leica Microsystems) allowed for precise targeting of these probes to desired locations, as well as continuous monitoring of electrode entry. The initial penetration through the AD, arachnoid, and pia was done at a higher speed (>100 µm/s), after which the penetration continued as slow speeds (2 µm/s). Following complete insertion, the probe was retracted slowly (2 µm/s) to relieve pressure without shifting the position of the probe relative to the cortex.

Electrical signals from the probe were collected at 30 kHz, digitized on a 64-channel headstage, and send to the recording controller (RHD Recording System, Intan Technologies). Action potential waveforms were extracted offline with Kilosort2^56,57^ with default settings (threshold = [10, 4], lambda = 10, AUC for splitting = 0.9) and manually sorted into single- and multi-unit clusters. To quantify the stability of our single unit recordings, we computed kernel density distribution estimates for several metrics^51–53^ (Figure S2E). Presence ratio reflects the proportion of a session during which a unit was present and firing action potentials. Maximum drift is the distance between the highest and lowest channels on which a unit was detected during a session. Isolation distance and d-prime quantify the separation of spike waveform clusters in principal component space. Single-unit clusters were further classified into broad- and narrow-spiking units based on their trough-to-peak waveform duration^32,58^. Units with waveform durations less than 350 µs were labelled as narrow-spiking, while units with waveform durations greater than 350 µs were labelled as broad-spiking. Units with a maximum waveform amplitude preceding the trough were classified as axonal spikes and excluded. Recordings were collected over the course of 17 sessions (8 in monkey M, 9 in monkey D). In total, 923 single units were recorded (461 in monkey M, 462 in monkey D). Only single units with a significant spatial receptive field (89.06%), as determined by a chi-squared test, were considered for subsequent analysis.

### Behavioral Control and Eye Tracking

Behavioral experiments were controlled with NIMH Monkeylogic^59^ in MATLAB. Eye position and pupil diameter were continuously sampled at 120 Hz (ETL-200, ISCAN Inc.) and sent to the behavioral control system. Stimuli were presented on a monitor (BenQ XL2411; 60 Hz refresh rate) positioned 57 cm from the monkey. Tolerance windows for fixation control were one degree of visual angle.

### Receptive Field Mapping

Receptive fields (RFs) were mapped with Gabor patch stimuli (2-4 cycles/deg, 0.5-1.5 deg Gaussian half-width, 100% luminance contrast) on a square grid spanning the visual quadrant of interest (lower right in monkey M, lower left in monkey D) while the subject maintained fixation. Grid spacing parameters were optimized for each session based on receptive field eccentricity and ranged from 0.25 – 1.0 degrees of visual angle (dva). A single Gabor was presented at one of six orientations (0, 30, 60, 90, 120, 150°) and at a grid location, both chosen at random, on each frame of stimulus presentation (60 Hz). Stimulus-evoked local field potential (LFP) power at each grid location on each recording channel was smoothed with a Gaussian kernel, and the peak location was defined as the RF center. Spatial RF maps for each channel were plotted as stacked contours for each shank to aid in visualization.

### Current Source Density Mapping

Current source density (CSD) mapping^31,32^ was used to identify laminar boundaries. While subjects held fixation, 100% contrast annular stimuli were flashed for 32ms, positioned over the center of the RF. The CSD was calculated as the second spatial derivative of the LFP. CSD traces were spatially smoothed with a Gaussian kernel (sigma = 140µm). The input layer was identified by an early current sink, representing feedforward input into layer IV. Channels above and below this sink were classified as superficial and deep respectively.

### Cued Saccade Task

During the task, subjects acquired and held fixation for a variable delay period (500-900 ms) prior to initiating a saccade in response to a target point appearing in the periphery. The simultaneous disappearance of the fixation point served as the go cue. After executing an accurate saccade, subjects then had to continue holding fixation at the target point for 500 ms to receive a reward. To prevent subjects from preemptively planning a saccade prior to the go cue, both the saccade target location and the delay period duration were pseudo-randomized. The target location was drawn from one of two possible locations, while the delay period duration was drawn from an exponential distribution. Targets were located 2.8 dva from the initial fixation point. Target locations were orthogonal to one another, and were each oriented 45 degrees to and equidistant from the fixation to receptive field axis. Only eye movements originating from < 0.75 dva of the initial fixation point, and terminating < 0.75 dva from the target were considered successful trials. While the subjects executed these eye movements, oriented Gabor stimuli were continuously presented on a 13 x 13 grid spanning the visual region of interest at 60 Hz. The grid was centered on a point 4 dva from fixation along the fixation-receptive field axis. On each frame of stimulus presentation, a single stimulus drawn from one of 6 random orientations (0, 30, 60, 90, 120, 150 degrees) was presented at a single grid location. Saccades were identified from eye-tracker data with a velocity-thresholding algorithm.^60,61^ On average, subjects performed 895 trials of the cued saccade task (minimum of 729 trials, maximum of 1029 trials).

### Continuous Receptive Field Mapping

Receptive fields were mapped for each single unit as a function of time. Spikes were binned for each unit in response to each stimulus flash in a time window 50 to 100ms after flash onset. Stimulus flashes were then binned (51 ms centered window slid from -400 to 400ms relative to saccade onset) and their corresponding spike counts were averaged. This procedure generated a 13 x 13 grid of spike counts at each location for each timepoint relative to saccade onset. Each timepoint was normalized such that the sum of all grid positions was equal to one to control for changes in firing rate. For visualization (Figure 2A only), this spatial grid was smoothed with a Gaussian kernel. The location of the current and future fields was determined by finding the stimulus position that elicited the maximum firing in the pre- and post-saccadic time periods, respectively. To compute sensitivity, the spike counts at the current and future fields were normalized to sum to one at each time point, such that the sensitivity reflects the relative proportion of firing in response to a stimulus presentation at the given field. Sensitivity analyses were done on the unsmoothed, spike count data. To determine the time at which each neural subpopulation first began to show remapping, we computed a bootstrapped 95% interval for the baseline sensitivity (-100 to -50 ms relative to saccade onset) at the future field. The first increase in sensitivity beyond these bounds was marked as the start of remapping for each bootstrapped population mean. Lastly, we determined the proportion of single units that showed clear remapping. Single units were considered to be remapping pre-saccadically if future field sensitivity exceeded (and remained above) current field sensitivity beginning at a timepoint before saccade onset. Only units with firing rates greater than 5 Hz were included in this proportion analysis (175 total units).

### Principal Component Analysis

A 13 x 13 grid of sensitivity was generated for each timepoint, as detailed above. Each timepoint was then vectorized to produce a 169-dimensional sample for each timepoint. All units from a given session on trials towards one of the two targets were fed into a PCA together, as they had overlapping current and future field locations in that condition. Each sample for a PCA thus reflects the 169-dimensional stimulus response space from one single unit at one timepoint. To average PCA results across sessions, values along the 1^st^ and 2^nd^ principal component axes were range normalized between 0 and 1 for each session. The time at which each subpopulation shows a significant change along the 1^st^ principal component axis was computed with bootstrapping, as described above for the sensitivity analysis.

### Tuning

To compute the tuning curves for each unit, spikes were binned in response to stimulus flashes of a given orientation at all positions within one of three epochs: pre-saccadic (-250 to -200 ms before saccade onset), saccade planning (-75 to -25 ms before saccade onset), and post-saccadic (150 to 200 ms after saccade onset). An orientation selectivity index and circular variance were calculated for each unit in each of the three epochs^62^. Each unit was fit with a Gaussian plus constant model function using the gaussfitn toolbox in MATLAB. Changes in orientation selectivity index, circular variance, firing rate, and half width at half height were quantified with the estimation stats toolbox^63^. Only units with a session-wide firing rate greater than 1 Hz were included in the tuning analysis.

## Notes

### Competing Interest Statement

The authors have declared no competing interest.

### Summary of Updates

Added new figures to further characterize the principal component analysis. Corrected a minor error in normalization procedures (does not affect findings, but reduces noise in analyses).

